# An ACE2 decamer viral trap as a durable intervention solution for current and future SARS-CoV

**DOI:** 10.1101/2023.07.10.548424

**Authors:** Hailong Guo, Bomsoo Cho, Paul R Hinton, Sijia He, Yongjun Yu, Ashwin Kumar Ramesh, Jwala Priyadarsini Sivaccumar, Zhiqiang Ku, Kristen Campo, Sarah Holland, Sameer Sachdeva, Christopher Mensch, Mohamed Dawod, Annalis Whitaker, Philip Eisenhauer, Allison Falcone, Rebekah Honce, Jason W. Botten, Stephen F Carroll, Bruce A Keyt, Andrew W Womack, William R Strohl, Kai Xu, Ningyan Zhang, Zhiqiang An, Sha Ha, John W Shiver, Tong-Ming Fu

## Abstract

The capacity of SARS-CoV-2 to evolve poses challenges to conventional prevention and treatment options such as vaccination and monoclonal antibodies, as they rely on viral receptor binding domain (RBD) sequences from previous strains. Additionally, animal CoVs, especially those of the SARS family, are now appreciated as a constant pandemic threat. We present here a new antiviral approach featuring inhalation delivery of a recombinant viral trap composed of ten copies of angiotensin-converting enzyme 2 (ACE2) fused to the IgM Fc. This ACE2 decamer viral trap is designed to inhibit SARS-CoV-2 entry function, regardless of viral RBD sequence variations as shown by its high neutralization potency against all known SARS-CoV-2 variants, including Omicron BQ.1, BQ.1.1, XBB.1 and XBB.1.5. In addition, it demonstrates potency against SARS-CoV-1, human NL63, as well as bat and pangolin CoVs. The multivalent trap is effective in both prophylactic and therapeutic settings since a single intranasal dosing confers protection in human ACE2 transgenic mice against viral challenges. Lastly, this molecule is stable at ambient temperature for more than twelve weeks and can sustain physical stress from aerosolization. These results demonstrate the potential of a decameric ACE2 viral trap as an inhalation solution for ACE2-dependent coronaviruses of current and future pandemic concerns.

## Introduction

Since its initial emergence into human populations in late 2019(*1, 2*), SARS-CoV-2 has been rapidly evolving into overlapping waves of several major variants including Alpha, Beta, Gamma, Delta, and Omicron, extending the pandemic over multiple seasons(*3*). Such remarkable dynamic evolution highlights the capability of this coronavirus (CoV) to adapt for efficient replication in its new host, and to effectively evade the host immunity. The SARS-CoV-2 viral mutation rate has been reported as 4-8 Χ 10^-4^ substitutions per site per year, which is only several fold lower than that of highly evolving influenza A virus(*4, 5*). A significant portion of these mutations have been identified in the receptor binding domain (RBD), a key structural component on the viral spike protein that binds to human angiotensin-converting enzyme 2 (ACE2) to initiate infection(*6*). Antibodies elicited by infection or vaccination have led to identification of hundreds of human monoclonal antibodies (mAbs)(*7*) that mostly target RBD for neutralization, and some have been developed and approved via emergency use authorization (EUA) for human use based on efficacy against the original wild-type (WT) Wuhan-Hu-1 strain, and early variants such as Delta(*8*). However, the emergence of Omicron and its sublineages, including the latest variants BQ.1, BQ.1.1 and XBB.1.5, has rendered nearly all mAbs, either approved for human use or in development, ineffective(*9–11*). These results indicate that therapeutics targeting antigenic epitopes on the RBD are not able to keep pace with CoV-2 viral evolution. Therefore, an alternative intervention strategy other than mAbs, with broad and durable efficacy, is urgently needed.

There are at least three human CoVs, (NL63, SARS-CoV-1 and SARS-CoV-2), that utilize ACE2 as the primary receptor for infection in human (*2, 12–14*). The specific sites that interact with the RBD of SARS-CoV-1 and CoV-2 have been mapped onto the N-terminal alpha helices on ACE2(*15, 16*), albeit with different contact residues. More strikingly, the RBD of NL63 interacts with a different structural footprint on ACE2 to that of the two SARS viruses (*17*). Thus, receptor binding motifs of coronavirus RBD can be diverse even for the same receptor, and this in turn explains why very few RBD-targeting antibodies for SARS-CoV-2 have been reported to cross neutralize other CoVs, or even SARS-CoV-1. In addition to SARS-CoV and NL63, newly emerged bat CoVs (e.g., WIV1 and BANAL-20), and pangolin CoVs (e.g., PCoV-GX and PCoV-GD), also use ACE2 for infection, and are thus identified as pandemic threats (*18–21*). Despite the vast difference in RBD sequences, all these viral isolates share a common functional attribute: they all depend on ACE2 for infection. Thus, it is possible to develop recombinant receptor decoys to trap viruses, as exemplified in the early work for hepatitis A virus, HIV-1 and rhinovirus (*22–24*). Recent studies demonstrate the feasibility of using soluble ACE2 or its IgG1 Fc fusion proteins for neutralizing SARS-CoV-2(*25–29*). However, these designs have the limitation of low neutralization potency, even with bivalent presentation on IgG1 Fc, commonly with two-to-three orders of magnitudes lower affinity to viral antigens as comparing to those of elite mAbs(*7*).

The IgM antibody format can significantly improve antiviral potency of IgG1-based mAbs due to the inherent multivalency effect(*30*). Here, we leveraged this property to construct IgM-based virus trapping molecules that present ten copies of ACE2. These ACE2 decamers, constructed on the IgM Fc frame that is composed of IgM Cµ3, Cµ4, tailpiece, and J chain (*31–33*), can be expressed and assembled efficiently in cultures. Furthermore, these molecules demonstrated over 10,000-fold improvement in RBD binding compared to ACE2 IgG1 fusions and can neutralize with single digit picomolar IC_50_ potency against all major SARS-CoV-2 variants, SARS-CoV-1, and recently isolated CoVs from bat and pangolin. A single intranasal administration of the lead ACE2 decamer in mice demonstrated prophylactic and therapeutic efficacy against WT and Omicron BA.2 challenge. In addition, the lead ACE2 decamer was aerosolized without any compromise of its biochemical and functional properties. These data highlight the medical potential of the IgM-based multimeric ACE2 as a virus trapping candidate that is potentially resilient to viral evolution and possesses a robust biochemical profile for development as an aerosol intervention for all coronaviruses that utilize ACE2 as receptor for infection.

## Results

### Multimerization of ACE2 ectodomain improves binding

Full length human ACE2 is composed of a signal peptide (SP) and a peptidase domain (PD), followed by a collectrin like domain (CLD), including a neck domain (ND), a transmembrane domain (TM) and intracellular domain (IC)(*34*). Since the ND (residue 616 to 768) mediates ACE2 dimerization, the ectodomain ACE2 can be expressed as a soluble dimer (dACE2, residues 18 to 740) with ND dimerization, or monomer ACE2 (mACE2, residue 18 to 615) without ND (Fig. 1A). Both mACE2 and dACE2 were fused to the IgM Fc frame via an IgG1 γ1 hinge linker (Fig. 1B). The mACE2-IgM and dACE2-IgM molecules were expressed with J chain in cell cultures (*29,30*) and assembled to form proteins with projected molecular weights of 985 and 1129 kDa, respectively. The IgG1 Fc versions of mACE2 and dACE2 were also expressed as controls (Fig. 1, B and C). The molecular integrities and authenticities were confirmed by SDS-PAGE and Western Blot (Fig. 1C and fig. S1). SEC analysis showed more than 95% homogeneity for all ACE2 molecules except dACE2-IgM, which was at 88%, displaying some aggregation possibly due to dimerization by ND (Fig. 1D). The results confirmed that monomeric or dimeric ACE2 can be efficiently expressed and assembled into IgM Fc.

**Fig. 1.**
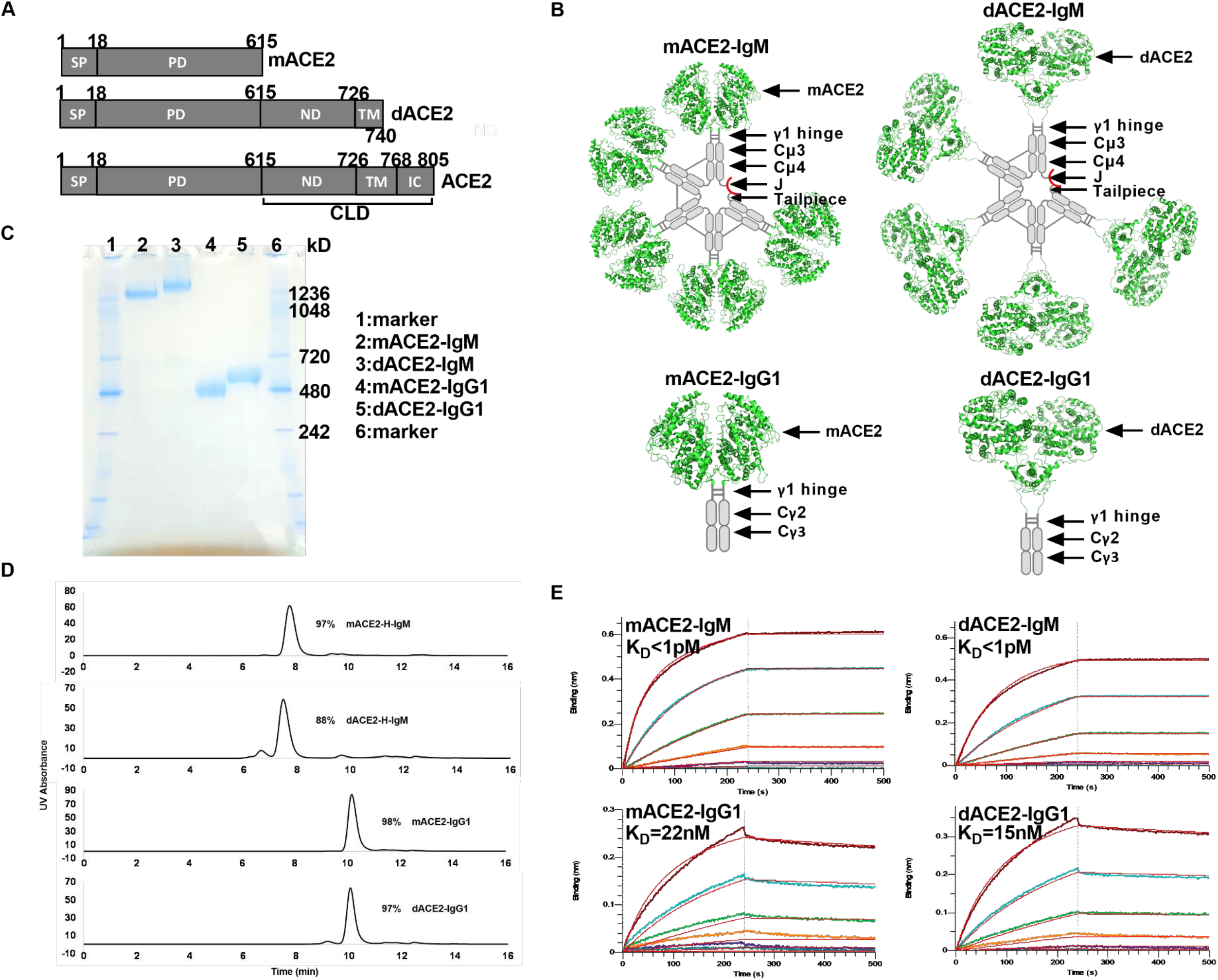
Design and expression of ACE2 decamer. (**A**) Illustration of the ACE2 truncations to design monomer or dimer ACE2 (mACE2 and dACE2). SP: signal peptide, PD: peptidase domain, ND: neck domain, TM: transmembrane domain, IC: intracellular domain, CLD: collectrin like domain. (**B**) Schematic diagrams of mACE2 and dACE2 in fusion with IgM Fc frame to form ACE2 decamer or IgG1 Fc as control using the available ACE2 structure (PDB: 6M1D). (**C**) Hybrid gel analysis of mACE2-IgM, dACE2-IgM, mACE2-IgG1, and dACE2-IgG1 expressed in fresh Expi293 culture supernatants after transient transfection, with Coomassie blue staining. (**D**) Size exclusion chromatography (SEC) analysis of purified mACE2-IgM, dACE2-IgM, mACE2-IgG1, and dACE2-IgG1. The purity estimate was based on the main peak. (**E**) Biolayer interferometry (BLI) analysis of Omicron BA.1 RBD binding by mACE2 and dACE2 decamer. The details of kinetic profiles are shown in table S1.

The binding kinetics of these molecules to Omicron BA.1 RBD were analyzed using bio-layer interferometry (BLI), and the multimerization by IgM Fc showed significant improvement of both on- and off-rates over the ACE2 constructs fused with IgG1 Fc (Fig. 1E and table S1). The *K*_D_ values of either mACE2-IgM or dACE2-IgM cannot be accurately assessed by these methods due to tight binding, but are estimated to be below 1 pM, representing more than a 10,000-fold increase over the corresponding IgG1-based constructs. These results indicate that decameric ACE2 displayed improved binding potential to the spike protein of SARS-CoV-2.

### Potent and broad viral neutralization

The antiviral biological functions of the generated mACE2-IgM decamers were first assessed in assays against lentivirus pseudotyped with the SARS-CoV-2 spike protein. Both mACE2-IgM and dACE2-IgM potently neutralized a panel of CoV2 variants including WT (D614G), Alpha, Beta, Gamma, Delta, and ten Omicron subvariants (BA.1, BA.2, BA.2.12.1, BA.2.75, BA.4/5, BQ.1, BQ.1.1, XBB.1 and XBB.1.5) in a dose dependent manner, with IC_50_ values ranging from 0.2–7.7 ng/mL (fig. S2A and table S2). In contrast, none of the ACE2-IgG1 constructs exhibited IC_50_ values ≤ 10 ng/mL and **>**50% of them had IC_50_ values above 100 ng/mL against corresponding pseudoviruses (table S2). The geometric mean IC_50_ against all SARS-CoV-2 pseudoviruses tested was 1.0 ng/mL (0.9 pM) for dACE2-IgM, and 1.9 ng/mL (1.9 pM) for mACE2-IgM with a statistical difference in a paired two-tailed *t*-test (p=0.030 for mass concentration) (Fig. 2A and fig. S2B). In the IgG1 Fc framework, dACE2-IgG1 also appeared to be several folds more potent with a geometric mean IC_50_ of 77.5 ng/mL (353.8 pM), as compared to that of mACE2-IgG1 at 372.0 ng/mL (1948.0 pM) with statistical significance (p=0.026 for mass concentration) (Fig 2A and fig. S2B). Both mACE2-IgM and dACE2-IgM were more potent than the corresponding IgG1 Fc fusion for all SARS-CoV-2 pseudoviruses with over 200-fold improvement on the geometric means for each pair (Fig. 2A and fig. S2B). These ACE2 decamers were also effective against SARS-CoV-1 Urbani pseudovirus(*35, 36*), with IC_50_ for the dACE2-IgM and mACE2-IgM at 2.4 ng/mL and 11.9 ng/mL, respectively (Fig 2B). Overall, the decameric mACE2 or dACE2 molecules demonstrated superb potency against pseudotyped SARS virus variants with single digit picomolar IC_50_ values, and both molecules were more potent in neutralizing various SARS pseudoviruses than their counterparts in the IgG1 Fc framework.

**Fig. 2.**
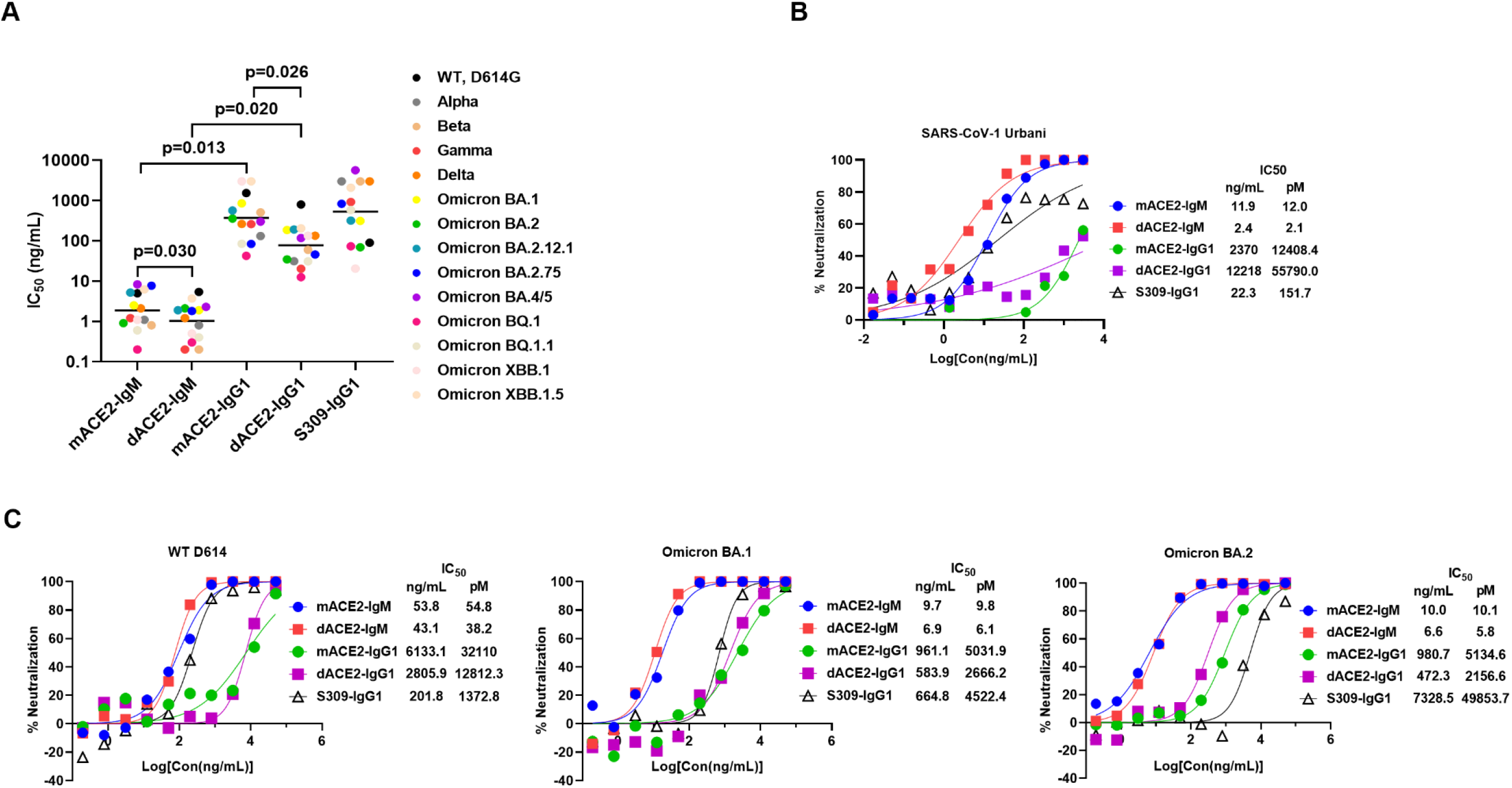
Virus neutralization of ACE2 decamer. (**A**) IC_50_ (ng/mL) of mACE2 and dACE2 decamers against a panel of pseudotyped SARS-CoV-2 variants. The panel includes major variants of Alpha, Beta, Gamma, Delta, and Omicron BA.1, BA.2, BA.4/5, BQ.1, BQ.1.1, XBB.1, and XBB.1.5, and each symbol represents one individual variant. The horizontal line indicates the geometric mean for the cluster. The IC_50_ values are listed in table S2. The statistical comparisons between indicated groups were conducted using two-tailed paired t-test. (**B**) Neutralization by mACE2 and dACE2 decamers against WT, Omicron BA.1 and BA.2 authentic virus. The IC_50_ (ng/mL) values represent means of at least two repeats. A recombinant IgG1 S309 was included as assay control.

The ACE2 fusions were further tested in culture against authentic viruses of WT WA1, Omicron BA.1, and BA.2. Both mACE2-IgM and dACE2-IgM decamers exhibited potent neutralization with IC_50_ values ranging from 7 to 54 ng/mL, with over 500-fold increase in potency for mACE2-IgM on a molar basis over mACE2-IgG1, and over 300-fold improvement for dACE2-IgM over dACE2-IgG1 (Fig. 2C). Similar to pseudovirus results (Fig. 2A), dACE2-IgM was slightly more potent than mACE2-IgM for authentic virus neutralization (Fig. 2C). However, because of the tendency of dACE2-IgM to aggregate (Fig. 1D), we selected mACE2-IgM for further evaluations.

### In vivo efficacy against SARS-CoV-2 challenge

The mACE2-IgM was administered intranasally into human ACE2 transgenic mice at different doses, either six hours before viral challenge for prophylaxis, or six hours after as therapy (Fig. 3A). In the prophylactic setting, after challenge with WT SARS-CoV-2 (WA1) virus intranasally at 10^3^ PFU, mice that received the irrelevant control IgM showed rapid and severe weight loss approaching 30% of the initial weights (Fig. 3B). In contrast, animals pretreated with either 0.5 or 5.0 mg/kg mACE2-IgM experienced only transient and mild weight loss of less than 7% during the 14-day monitoring period (Fig. 3B). The viral titers measured in the lung tissues collected at day 3 post challenge were significantly lower in mice treated with mACE2-IgM than the IgM control (p=0.0061) (Fig. 3C). Histopathological examination by H&E staining revealed massive hyperemia, thickened, or broken alveolar epithelial walls, and marked peribronchial and perivascular immune cell infiltrations in the lungs of control mice, but not in the two groups that received mACE2-IgM (fig. S3A).

**Fig. 3.**
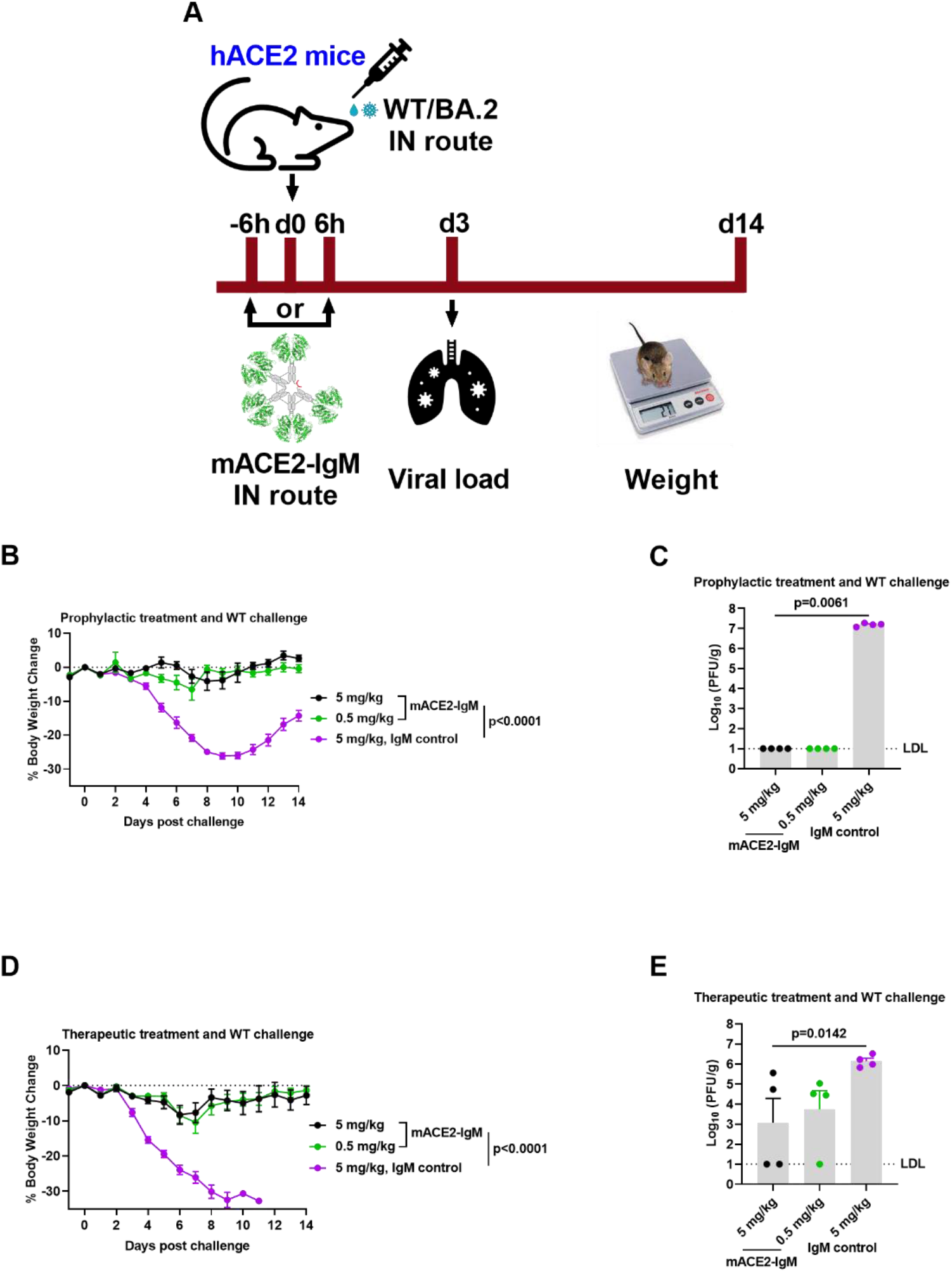
In vivo protection of ACE2 decamer. (**A**) Schematic illustration of test article treatment with different doses in K18 hACE2 delivered intranasally into transgenic mice (12 per group) 6h before or after intranasal challenge with 1000 PFU of WT WA1 or 7.0X10^3^ TCID_50_ of Omicron BA.2 virus. (**B**) Weight change and (**C**) viral titer of lungs of animals treated intranasally with mACE2-IgM or control 6h before intranasal challenge with WT WA1 virus. (**D**) Weight change and (**E**) viral titer of lungs of animals treated intranasally with mACE2-IgM or control 6h after intranasal challenge with WT WA1 virus. The data are means ± SEM for weight change and viral load in each group. Two-way ANOVA (mixed-effects model) and nonparametric one-way ANOVA were used for comparisons of weight change and lung viral titer of different groups, respectively. LDL: lower detection limit.

In the therapeutic setting, mice given the control IgM treatment showed rapid and severe weight loss leading to 100% mortality (Fig. 3D) while animals receiving either 0.5 or 5.0 mg/kg mACE2-IgM experienced only mild weight loss no more than 10% mortality (Fig. 3D). The viral titers measured in the lung tissues collected at day 3 post challenge were also significantly lower in the treatment groups than the control (p=0.0142), with no virus detected in one mouse out of four in the 0.5 mg/kg group and two of four in the 5 mg/kg group (Fig. 3E). Further, the histopathology study of the lungs of control mice on day 3 showed severe edema, distorted alveoli, and massive peribronchial and perivascular immune cell infiltrations, whereas mice in the treatment groups showed minimal tissue pathology (fig. S3B).

We also conducted similar challenge studies with an Omicron BA.2 isolate at 7.0X10^3^ TCID_50_ in hACE2 transgenic mice, either treated with mACE2-IgM prophylactically or therapeutically, with a dose ranging from 0.625 to 5.0 mg/kg. In both settings, we observed clear protection against weight loss when dosed with mACE2-IgM, at all dose levels, as compared to the control group (fig. S4A and S4C). The mice receiving mACE2-IgM survived better than the controls (p=0.096 for prophylactic treatment and p=0.005 for therapeutic treatment). In addition, the viral titers in lung tissues collected at day 4 post challenge showed reduction for the treatment groups as compared to the control groups (fig. S4B and S4D). These preclinical data confirmed the interventional benefits of mACE2-IgM in an animal model and suggest that intranasal administration of the decameric ACE2 is suitable for further development.

### Aerosolization and developability of mACE2-IgM

Drug delivery via inhalation is a promising and non-invasive administration for respiratory disease (*37, 38*). However, inhalation delivery commonly requires aerosolization, propelled by mechanical vibration and pressurization, which is a well**-**identified physical stress to large protein molecules. Formulation optimization was conducted to stabilize the multimeric ACE2 during atomization and nebulization, and a lead formulation was demonstrated to enable mACE2-IgM aerosolization with 100% protein recovery for both atomization and nebulization and without any detectable increase in high molecular weight (HMW) species, indicating strong stability against shear stress (Fig 4A and table S3). Importantly, atomization and nebulization had no impact on mACE2-IgM binding as indicated by ELISA (table S3). Taken together, mACE2-IgM was successfully aerosolized while maintaining structural and functional integrity, paving the way for its development as an aerosolized biologic.

**Fig. 4.**
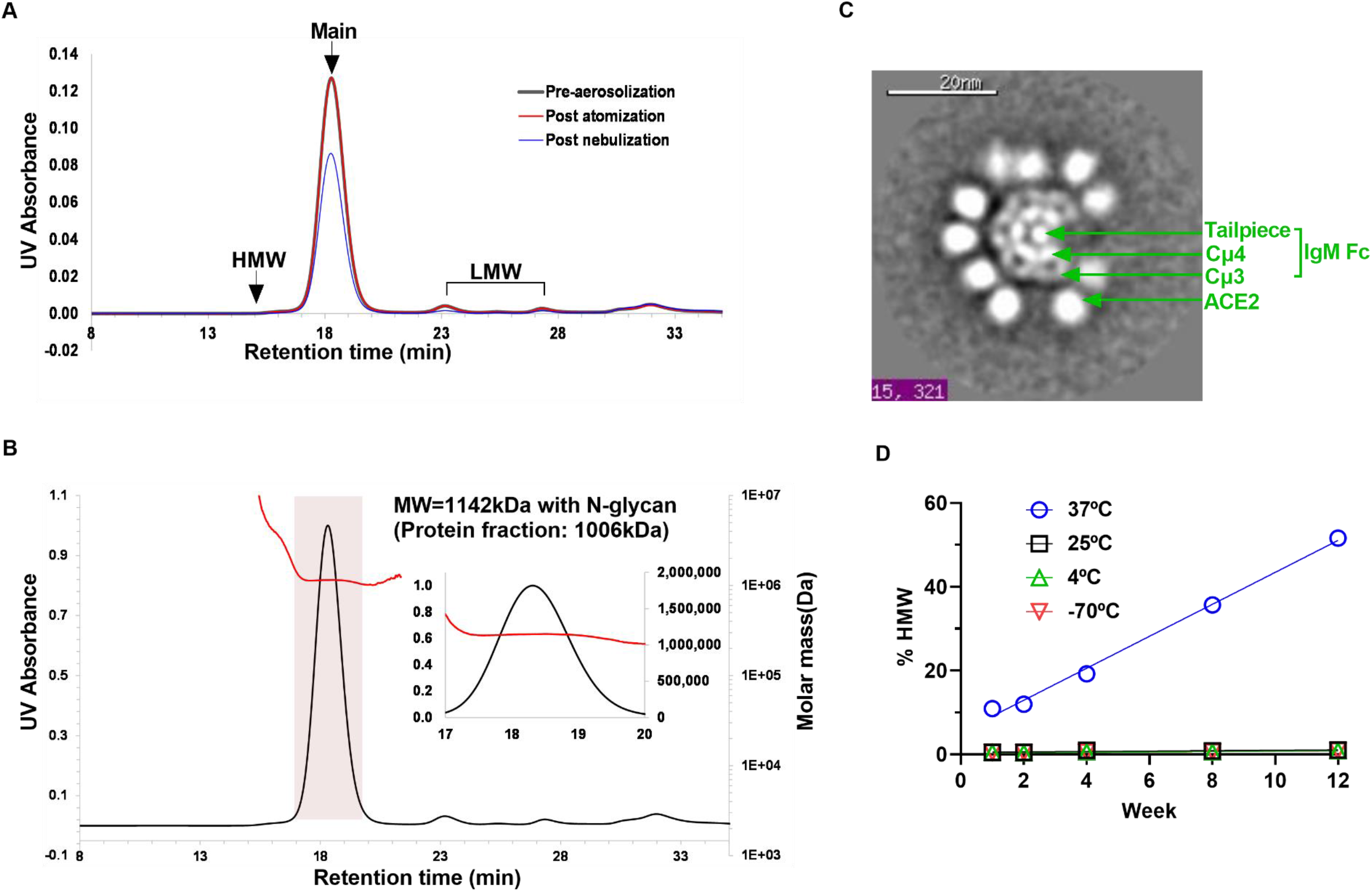
Aerosolization and developability assessment of ACE2 decamer. (**A**) SEC chromatograms of mACE2-IgM pre- and post-aerosolization either by atomization using MAD 300 or by nebulization using Aerogen Solo. The HMW denotes aggregation species while the LMW species were protein impurities in this specific batch. (**B**) Homogeneity analysis of mACE2-IgM by SEC-MALS. The black line is the UV trace, and the red line is the calculated molar mass. The insert panel is a zoom-in view of the main peak with the left axis showing the calculated molar mass in a linear scale. (**C**) Example of the TEM 2D class averages of mACE2-IgM with the densities surrounding the IgM Fc frame representing 10 copies of ACE2 domains. The first number at the bottom states the class number and the second number indicates the number of particles contributing to that class. (**D**) Stability of mACE2-IgM upon storage at different temperatures including −70°C, 4°C, 25°C, and 37°C detected by SEC-UV.

Although tremendous knowledge has been gained regarding IgGs and IgG-fusion biologics development in the past decades, IgM-multimerized biologics have been less studied. Thus, we conducted developability characterizations on the ACE2 decamer to address potential risks such as stability, uncontrolled heterogeneity, or unexpected modification. First, we selected the monomer version of ACE2 which lacks the whole CLD including neck region, and this in turn reduces aggregation liability (Fig. 1A, D). Second, we included the J chain in the assembly, which locks the molecules in the homogeneous decamer form. Size exclusion chromatography and multi-angle light scattering showed the main population of molar mass (MW) of 1006 kDa, consistent with the theoretical MW of 985 kDa for a decamer (Fig. 4B). The measured MW across the entire main population was constant, indicating that mACE2-IgM formed highly homogeneous decamers, not a mixture with 12-mers (Fig. 4B). Third, the mACE2-IgM molecule was expressed at high levels in both Expi293 and CHO mammalian cultures and assembled in decamer form as confirmed by negative stain transmission electron microscopy (TEM) (Fig. 4C). Most importantly, mACE2-IgM did not form aggregates at ambient or lower temperature up to 12 weeks, in contrast to incubation at 37°C, which showed steady aggregation after 2 weeks of storage (Fig. 4D). Furthermore, mACE2-IgM maintained reactivity to RBD after storage at 25°C for at least 12 weeks as tested by ELISA. The ambient temperature stability is indicative of the robust structural and biochemical properties of the candidate. These profiles collectively indicate that mACE2-IgM can be developed as an aerosolized biologic.

### Neutralization against human NL63 and emerging animal CoVs

All known human CoVs including common-cold CoVs such as alpha CoVs NL-63 and 229E, and severe disease-causing CoVs such as SARS-CoV-1 and SARS-CoV-2, originated from animal reservoirs (*2, 39*). NL63 can use ACE2 for infection that can also lead to severe diseases in infants and patients with underlying immune disorders (*17, 40–43*). The bat CoVs WIV1 and BANAL-20, and the Pangolin PCoV-GX and PCoV-GD are recently discovered animal CoVs that are closely related to SARS-CoV-1 or SARS-CoV-2 (Fig. 5A) with the capability of using ACE2 as receptor to infect humans (*18–21*). To demonstrate the potential of mACE2-IgM as an intervention against pandemic threats of these animal CoVs and severe infection of NL63, we constructed recombinant RBD and pseudovirus panels corresponding to these CoVs. The kinetic study by BLI showed that mACE2-IgM can bind to each recombinant RBD with high affinity despite poor homology with WT SARS-CoV-2 for some of these RBDs (Fig. 5B and fig. S5). These data indicate the potential antiviral activity of mACE2-IgM for the corresponding viruses. Indeed, neutralization studies demonstrated potent antiviral activity of mACE2-IgM against pseudoviruses of WIV1, BANAL-20, PCoV-GD, PCoV-GX and NL63 with IC_50_ values ranging between 4-20 ng/mL (Fig. 5C). These results reveal that mACE2-IgM could be an excellent candidate for treating severe diseases caused by NL63 and other emerging ACE2-dependent animal CoVs with pandemic potential.

**Fig. 5.**
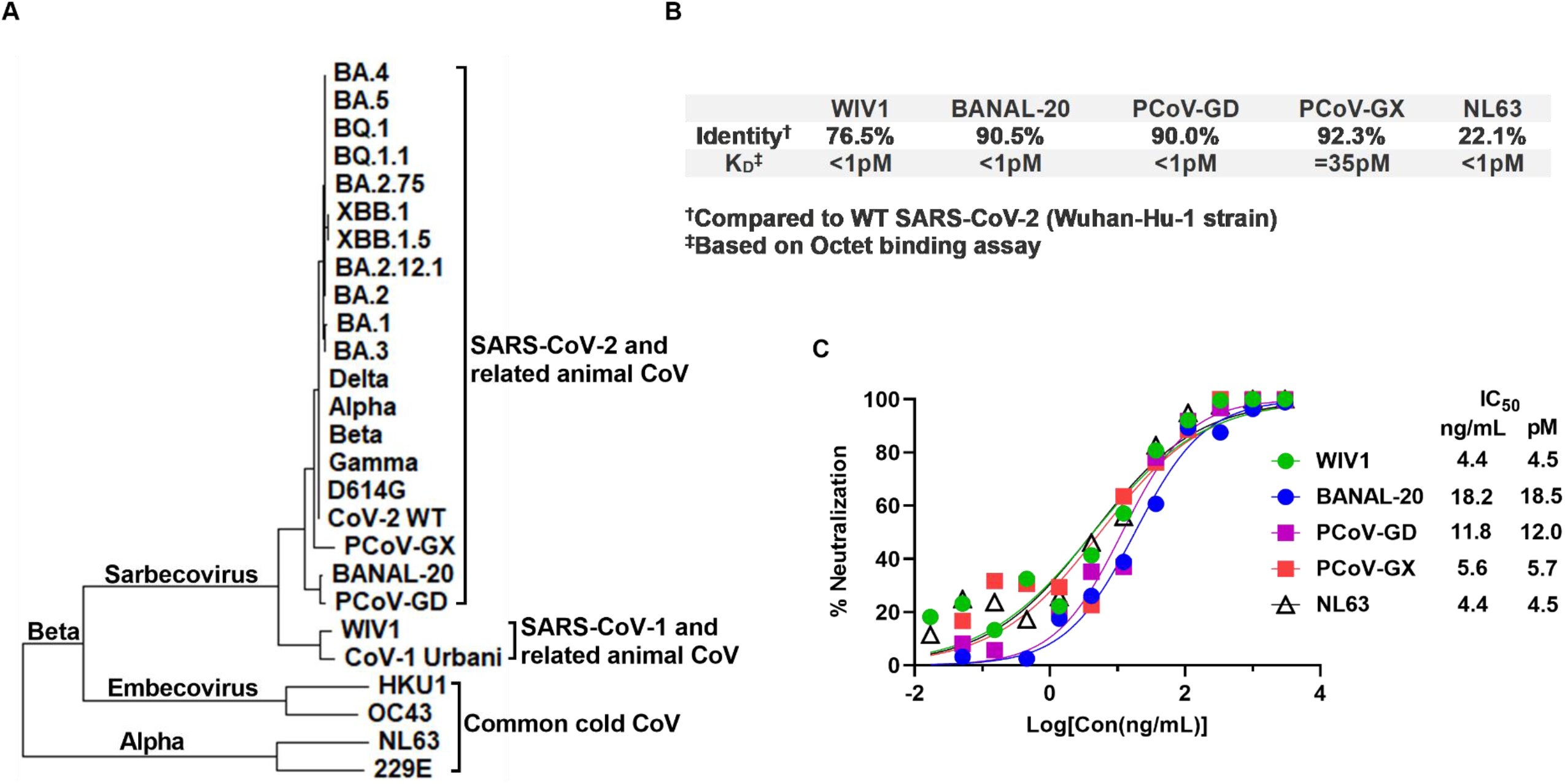
Neutralization of emerging animal CoV and NL63 by mACE2-IgM decamer. (**A**) Phylogenetic tree of spike proteins of alpha and beta CoVs including major variants of SARS-CoV-2, SARS-CoV-1, and the closely related animal CoVs, and four human common cold CoVs. (**B**) Sequence identity of the spike of WIV-1, BANAL-20, PCoV-GD, PCoV-GX, and NL63 viruses with WT SARS-CoV-2 and the K_D_ of mACE2-IgM binding derived from BLI assay. (**C**) Pseudovirus neutralization against WIV-1, BANAL-20, PCoV-GD, PCoV-GX, and NL63 by mACE2-IgM. IC_50_ values were calculated based on four parameter curve fitting.

## Discussion

The pandemic experience of SARS-CoV-2 provides important learnings about CoVs for the scientific community. First, zoonotic CoV poses pandemic threat as dangerous as influenza, if not more so. For instance, SARS-CoV-2 has demonstrated the ability to rapidly gain infectiousness through its evolution, with the latest Omicron variants reaching an average *Ro* (denoted for reproduction number) of 10, far beyond that of influenza virus of 2 and approaching the rate of measles virus of 12-18, one of the most contagious viruses ever discovered(*44–46*). Second, literature and our data here showed existence of animal isolates of beta-CoV lineages that can use the same ACE2 for infecting human cells (*18, 19*). Also of note is how efficiently SARS-CoV-2 has adapted in new host species via only one (N501Y)(*47*) or two (Q493K and Q498H/Y) (*48, 49*) mutations in the RBD domain to bind better to new host ACE2 and replicate more productively. These examples illustrate an ancestral trait of CoVs, as their RBDs are highly plastic and can evolve readily for replication in new hosts (*50*). Third, dynamic viral evolution observed in SARS-CoV-2 has highlighted challenges to the development of durable and broad vaccines or therapeutic antibodies. Lastly, the primary SARS-CoV-2 replication at the respiratory mucosal epithelium is less effectively controlled by IgG antibodies in circulation, as only 1-2% of parenterally administered IgG antibody is detectable in the nasal lining fluid (*51*). The low concentrations of IgG antibodies on the upper respiratory mucosal surfaces, coupled with the high viral mutation rate, create a perfect milieu for SARS-CoV-2 escape mutants to arise, nullifying antibody responses. This scenario may contribute to accelerated viral evolution, as seen with SARS-CoV2 and 229E (*52*).

Here we present a new antiviral concept that can directly address these challenges by first targeting viral entry function using a viral trap enabled by IgM-multimerization and second by using inhalation delivery to the site of active viral infection. Our data, exemplified by the mACE2-IgM, demonstrate the promise of such an approach in preclinical settings and developability of such large molecules as an aerosolized direct antiviral. Our approach focuses on blocking viral entry by viral trapping, a sharp departure from the conventional wisdom established by antiviral mAbs or vaccines that rely on viral antigen sequences (*7*). Thus, in contrast to mAbs, decameric viral traps should maintain their effectiveness, and no longer be subject to viral evolution and antigenic diversity commonly seen in respiratory viruses. Although there is a theoretical possibility that the coronavirus could adapt to an alternative receptor for infection or become ACE2-independent, to our knowledge such a concern has not been experimentally validated.

Monomeric soluble ACE2 exhibits low potency in virus neutralization assays (*29, 53–55*), as the reported IC_50_ is about three orders of magnitude lower than those of elite RBD-specific antibodies(*40*). One solution is to engineer ACE2 to improve its affinity specifically to SARS-CoV-2(*25–28*). However, this approach deviates from native ACE2 structure and potentially limits the efficacy against future viral evolution. Another solution is to improve potency through valency, and ACE2 has been engineered through dimerization with IgG1 Fc, trimerization with T4-foldon, and tetramerization with engineered IgG1 Fc (*41*). Recently, one report also has shown a construct with twelve copies of ACE2 with an IgM framework without J-chain (*59*). These efforts showed incremental improvement on antiviral potency, consistent with our results. The key difference between our approach versus those reported is in engineering details, as we focused on developable IgM framework fusions composed of only Cμ3 and Cμ4 with tail piece and J chain, the key structural components for efficient IgM assembly into pentameric structures, which appear similar to native IgMs in EM (Fig. 4C) (*60*). In addition, ACE2 orientation and flexibility were improved with the incorporation of an engineered IgG1 hinge linker. These features plus formulation optimization greatly improved developability of the ACE2 decamer.

Animal CoVs, especially those of sarbecovirus clades of beta genus, have great potential to infect humans via ACE2 receptor as evidenced by the SARS-CoV-1 (2003) and SARS-CoV-2 (2019) outbreaks. CoV virus in other genus, such as NL63 of alpha genus, can also utilize ACE2 receptor for infection in humans (*40–43*). In addition, the SARS-CoV-2 virus has spread back to animal species, domesticated or wild, establishing new animal reservoirs that can foster new viral mutations and reinfect humans. This concern has been best illustrated by the recent findings that, in white-tailed deer populations, reverse zoonotic SARS-CoV-2 developed distinct mutational patterns and were transmitted back to human populations (*61–65*). Therefore, threats of CoV in animal reservoirs capable of transmitting to humans can be significant. Although mACE2-IgM is currently being developed as a candidate for intervention against COVID-19, based on our in vitro data it should also be effective against any human or zoonotic CoV that depends on ACE2 for infection.

In summary, we have demonstrated the feasibility of an IgM-enabled virus trapping strategy, here as an ACE2 decamer to block viral infection through ACE2. This molecule confers potent and broad neutralization against SARS-CoV-2 viral variants and selected animal CoVs and provides protection against SARS-CoV-2 challenge in mice. Inhalation delivery can directly deposit the decameric ACE2 to the mucosal site (*37, 38*), effectively blocking SARS-CoV-2 replication and further evolution, and potentially reducing its transmission at population levels. The molecule has shown robust biochemical and biophysical profiles suitable for further development as an aerosolized antiviral agent. This virus trapping molecule can be developed as a durable intervention for all CoV viruses that depend on ACE2 for infection.

## Methods

### Protein expression and purification

Human ACE2 fusion proteins were obtained from transient transfection using the Expi293^TM^ system (Thermo Fisher Scientific). All plasmids used in this study were generated by ATUM Bio and transient transfection was carried out following the manufacturer’s protocol for the ExpiFectamine^TM^ 293 transfection kit (Thermo Fisher Scientific). Four days after transfection, the supernatant was collected after centrifugation of the culture and filtered through a 0.2 µm filter to remove cells and debris. All purifications were performed using mixed-mode and anion-exchange chromatography as described previously(*30*).

### Hybrid gel electrophoresis and Western blot

Samples were mixed with NuPAGE™ LDS Sample Buffer (Thermo Fisher Scientific) and were analyzed on NativePAGE 3-12% Bis-Tris Gel (Thermo Fisher Scientific). The gels were run at 50V for 20 min and 100V for an additional 2 hrs in the Tris-acetate SDS running buffer (Thermo Fisher Scientific). For Western blot, samples were mixed with LDS Sample Buffer and reducing agent, boiled, and loaded on NuPAGE 4-12% Bis-Tris Gel (Thermo Fisher Scientific). The gels were stained with the InstantBlue Coomassie (Abcam) for 10min before analyzing with ChemiDoc™ Touch Imaging System (Bio-Rad). J chain Western blots were performed using iBlot^TM^ 2 Dry Blotting System (Thermo Fisher Scientific) and iBind^TM^ Flex Western system (Thermo Fisher Scientific) following the manufacturer’s protocol. Antibodies: Rabbit anti-J Chain (SP105, 1:100, Thermo Fisher Scientific), Goat anti-Rabbit IgG1, AP conjugated (1:1,000, Thermo Fisher Scientific), and 1-Step^TM^ NBT/BCIP (Thermo Fisher Scientific) was used for detection.

### Size exclusion chromatography (SEC) coupled to ultraviolet (UV) or multi-angle light scattering (MALS)

SEC-UV method consisted of an Agilent 1260 HPLC with a UV-Vis detector. An XBridge Protein BEH SEC Analytical Column (Waters, 450Å, 3.5 µm, 7.8 mm X 300 mm) was used for the separation. An XBridge Protein BEH SEC Guard Column (Waters, 450Å, 3.5 µm, 7.8 mm X 30 mm) was attached before the analytical column as a protection. The column was kept at 30°C. The UV detection was set at 220 nm. The flow rate was 1 mL/min. The SEC-MALS method consisted of an Agilent 1260 HPLC with UV, DAWN (Wyatt) MALS, and Optilab (Wyatt) relative refractive interferometer (RI). A Tosoh TSKgel G4000SWXL column (P/N 08542, 8 µm, 7.8 x 300 mm) was used for the separation. The flow rate was 0.4 mL/min. The column and mobile phase were held at room temperature. Data were analyzed using ASTRA software. Peak alignment and band broadening correction between UV, MALS, and RI detectors were performed. The dn/dc of 0.185 was used for the protein fraction and 0.147 for the glycan modifier.

### Negative stain transmission electron microscopy (TEM)

Negative stain TEM was conducted at NanoImaging Services (San Diego, CA). Briefly, mACE2-IgM at 0.045 mg/mL was applied to a copper mesh grid that was plasma-cleaned using a Gatan Solarus (Pleasanton, California), negatively stained with a 1% uranyl formate solution, blotted and air-dried. TEM was performed using an FEI Tecnai T12 TEM (serial number D1100), operating at 120keV with an FEI Eagle 4k x 4K CCD camera. Negative stain grids were transferred into the TEM using a room temperature stage. Images of each grid were acquired at multiple scales to assess the overall distribution of the specimen. After identifying potentially suitable target areas for imaging at lower magnifications, high magnification images were acquired at nominal magnifications of 67,000x (0.16 nm/pixel) and 110,000x (0.10 nm/pixel). TEM particles were identified in the high magnification images and subjected to two rounds of alignment to obtain class averages using the XMIPP processing package(*66*).

### Biolayer Interferometry

Biolayer interferometry (BLI) was carried out using Octet RH16 (Sartorius). Recombinant SARS-CoV-2 B.1.1.529 (Omicron BA.1) Spike RBD-His protein (Sino Biological) was diluted with the 10X Kinetic Buffer (Sartorius) to 0.5 µg/mL and immobilized to Octet® HIS1K Biosensors. The labeled biosensors were exposed to various ACE2 fusion proteins at concentrations ranging from 0 to 66.7 µg/mL in the 10X Kinetic buffer during the association period, then returned to the baseline well containing the 10X Kinetic Buffer during the dissociation period. The binding protocol was as follows: equilibrate sensors in baseline wells, 60 seconds; load RBD-His, 180 seconds; establish baseline in baseline wells, 60 seconds; association with analyte, 240 seconds; dissociation in baseline wells, 600 seconds. Sensors were regenerated with 10 mM glycine pH 3.0 between analytes. Raw data were fit to a bivalent model using Octet® Analysis Studio 12.2.2.26 software with global fitting.

### ELISA assay

384 well ELISA plates were coated with 20µl of RBD (SARS-CoV-2 Omicron BA.1, ACROBiosystems) at 1µg/mL in PBS at 4°C overnight. After standard washing and blocking, 20µl of diluted ACE2 fusion molecules and other control antibodies were added to each well. After one hour incubation at room temperature, plates were washed and incubated with HRP conjugated Goat anti-human IgG1 Fc fragment (A80-104P, BETHYL) for IgG1 and Goat anti-human IgM antibody (A80-100P, BETHYL) for IgM for 30 minutes at room temperature. SuperSignal^TM^ ELISA Pico Chemiluminescent Substrate (Thermo Scientific) was used for detection using the Microplate Reader (Perkin Elmer). EC_50_ values were calculated using four parameter nonlinear regression fitting (GraphPad Prism 9.2.0).

### Pseudovirus generation and neutralization

GFP lentiviral reporter virus particles (RVPs) pseudotyped with the spike protein of WT and all variants of SARS-CoV-2 and SARS-CoV-1 used in this study were from Integral Molecular (Philadelphia, PA). The pseudovirus containing the spike protein of human NL63 (GenBank: AAS58177.1), Bat SARS-like coronavirus WIV1 (GenBank: AGZ48828.1), Bat coronavirus BANAL-20-236 (GenBank: UAY13253.1), Pangolin coronavirus PCoV-GD (GenBank: QLR06867.1), Pangolin coronavirus PCoV-GX (GenBank: QVT76606.1) were prepared using the same lentiviral vector system. Clonal 293T-hsACE2 cells (Integral Molecular, Cat# C-HA102) were maintained in cell culture medium including DMEM, 10% FBS, 10 mM HEPES, 100 U/mL of Penicillin-Streptomycin and 0.5 μg/mL of Puromycin. For neutralization assays, 293T-hsACE2 cells were seeded at 1.5 × 10^4^ cells/well with 100 μL cell culture medium in 96-well plates. ACE2 molecules were prepared with 3-fold serial dilutions starting at 12 μg/mL (3 μg/mL to be the final concentration) and mixed with equal volume of 40000TU/ml GFP RVPs (Integral Molecular), followed by 1 hour incubation at 37°C. After the incubation, 100 μL of mixture was added to 100 μL cells in each well. The plate was incubated in a 5% CO2, 37°C incubator for 72 hours. The GFP signal was acquired by BioTek Cytation 7 imaging reader, and the virus entry events were counted by Gen5 software. IC_50_ values were calculated using a non-linear regression curve fit ([inhibitor] versus normalized response–variable slope) by GraphPad Prism 9.2.0 as described(*67*).

### Authentic virus neutralization

All live virus neutralization assays using SARS-CoV-2 were conducted at the University of Vermont BSL-3 facility under an approved Institutional Biosafety protocol. SARS-CoV-2 strain 2019-nCoV/USA_USA WA1/2020 (WA1) was generously provided by Kenneth Plante and the World Reference Center for Emerging Viruses and Arboviruses (WRCEVA) at the University of Texas Medical Branch. SARS-CoV-2 strain Omicron subvariant BA.1 and BA.2 were obtained through BEI Resources, NIAID, NIH. All SARS-CoV-2 strains were propagated in Vero E6 cells. The microneutralization assay itself was done with low passage Vero E6 cells. Each antibody sample was diluted in 50 µL of cDMEM. Antibody concentrations were tested in serial 1:4 dilutions. Diluted antibody samples were mixed with 50μL of cDMEM containing 175 focus forming units of WA1, or Omicron viruses and then incubated for 45 minutes at 37°C. To inoculate, the media from confluent Vero E6 cell monolayers in 96-well white polystyrene microplates (07-200-628, Thermo Fisher Scientific) was removed and 50 µL of each antibody-virus mixture was inoculated onto the cells in duplicate technical replicates and incubated at 37°C in a 5% CO2 incubator for 30 minutes, after which the wells were overlaid with 150μL of warmed 1.2% methylcellulose in cDMEM and incubated at 37°C in a 5% CO2 incubator for 24 hr before infected cells were fixed in 4% formaldehyde in 3X phosphate buffered saline (PBS). Cells were permeabilized with 0.1% 100X Triton in 1X PBS for 15 minutes at room temperature and then incubated with a primary, cross-reactive rabbit anti-SARS-CoV N monoclonal antibody (40143-R001, Sino Biological) (1:20,000) for 45 minutes at 37°C followed by a peroxidase-labeled goat anti-rabbit antibody (5220-0336, SeraCare) (1:2,000) for 30 minutes at 37°C and then the peroxidase substrate (5510-0030, SeraCare) was added to develop blue viral foci. Images of the wells were captured using the CTL Immunospot S6 Universal Analyzer, and viral foci were then quantified, with foci counts normalized to virus only control wells on each plate. IC_50_ determinations were made using a non-linear regression curve fit ([inhibitor] versus normalized response – variable slope) by GraphPad Prism 9.2.0 described previously(*67*).

### Sequences and phylogenetic tree analysis

Spike protein sequences for SARS-CoV-2 WT and variants were retrieved from GISAID database with the following ID: EPI ISL 402124 (Wuhan-Hu-1/WT), EPI ISL 14380457 (B.1/D614G), EPI ISL 16705748 (B.1.1.7/Alpha), EPI ISL 14716761 (B.1.351/Beta), EPI ISL 12625231 (P.1/Gamma), EPI ISL 15515853 (B.1.617.2/Delta), EPI ISL 13719390 (Omicron BA.1), EPI ISL 16803258 (Omicron BA.1.1), EPI ISL 9027794 (Omicron BA.2), EPI ISL 14367842 (Omicron BA.2.12.1), EPI ISL 14371488 (Omicron BA.2.75), EPI ISL 13265244 (Omicron BA.3), EPI ISL 16543330 (Omicron BA.4), EPI ISL 12568880 (Omicron BA.5), EPI ISL 16763343 (Omicron BQ.1), EPI ISL 16818999 (Omicron BQ.1.1), EPI ISL 16817457 (Omicron XBB.1), and EPI ISL 16818774 (Omicron XBB.1.5). The other sequences were derived from NCBI GenBank with the following ID: AAP13441 (SARS-CoV-1 Urbani), AGZ48828.1 (Bat SARS-like coronavirus WIV1), UAY13253.1 (Bat coronavirus BANAL-20), QVT76606.1 (Pangolin coronavirus PCoV-GX), QLR06867.1 (Pangolin coronavirus PCoV-GD), ABD96187.1 (human coronavirus HKU1), AAR01015.1 (human coronavirus OC43), AAS58177.1 (human coronavirus NL63), NP_073551.1 (human coronavirus 229E). These sequences were aligned with MUSCLE and the phylogenetic tree was constructed using using the Maximum Likelihood method and JTT matrix-based model with 1000 bootstrap replicates in MEGA_11.0.10.

### Animal and NL63 CoV RBD generation and Octet binding

RBD proteins of WIV1 (AGZ48828.1), BANAL-20 (UAY13253.1), PCoV-GX (QVT76606.1), PCoV-GD (QLR06867.1), and NL63 (AAS58177.1) containing Fc-tag were constructed as described previously(*30, 68*). Briefly, gene fragments of different RBDs were fused with human IgG1 Fc fragment and inserted into linearized expression vectors. Stellar™ Competent Cells (Takara Cat#: 636763) were used to generate clones of the vector containing RBD genes of interest. Colonies were selected and sequenced to confirm the presence of RBD gene of interest. Plasmids were expressed using an Expi293F (GIBCO, cat#100044202) cell-based expression system and RBD proteins were purified using Protein A affinity resin (Genscript Cat#: L00210). Protein purities were assessed by SDS-PAGE. The evaluation of the avidity of mACE2-IgM to these RBD proteins was performed at room temperature on the ForteBio Octet RED96 system. Fc-tagged RBD proteins (20 μg/mL) were captured on the protein A biosensor followed by 30 s of baseline run in 1x kinetics buffer (Sartorius Cat#: 18-1105). The biosensors were blocked with a control Fc protein (Jackson ImmunoResearch Cat#: 009-000-008) at 150 μg/mL for 200 s to occupy the free protein A. The sensors were dipped in threefold serially diluted mACE2-IgM (0.12 nM to 90 nM) for 300 s to record association kinetics. To record the dissociation kinetics, the sensors were dipped into kinetics buffer for 1800 s. ForteBio Octet data analysis software with 1: 1 model was used to fit the *K*_D_ data using the global fitting method.

### Animal studies

WT virus challenge studies were conducted at the University of Vermont ABSL-3 facility. All procedures (PROTO202000187) were approved by the University of Vermont Animal Care and Use Committee (IACUC) and complied with the Guide for the Care and Use of Laboratory Animals. 12-week-old male K18-hACE2 transgenic mice (Jackson Laboratory) were housed in HEPA-filtered, negative pressure, vented isolation containers and maintained at 12 h light-dark cycles and ambient temperature and humidity with food and water provided *ad libitum* for the duration of the experiment. For treatment, mice were lightly anesthetized with 3% inhaled isoflurane and instilled with indicated dosage (5 mg/kg or 0.5 mg/kg) of mACE2-IgM or controls via the intranasal route. For the prophylactic study group, drug instillation occurred approximately 6 hours pre-infection while the therapeutic study group was dosed 6 hours post-infection. Intranasal infection occurred under light anesthesia, with mice instilled with 10^3^ PFUs SARS-CoV-2 (WA1 strain) in 50 μL PBS. Mice were monitored daily for weight and clinical signs of infection. At greater than 30% body weight loss from baseline or clinical signs of greater than 3, mice were humanely euthanized following AVMA guidelines. Clinical signs were agreed upon by at least two researchers and scored as follows: 0=no observable signs, 1=active, squinting, hunching or scruffy appearance; 2=squinting and hunching, active only when stimulated; 3=excessive hunching, squinting, not active when stimulated; 4=hind-limb paralysis, shivering, rapid breathing, moribund and 5=death. On day 3 post-infection, mice were euthanized via CO_2_ inhalation and thoracotomy prior to removal of tissue for downstream assays. A 20 mg section of the right caudal lobe was dissected from each subject and immediately frozen at −80°C for titer determination via plaque assay with the remaining lung inflated, excised, and fixed with 10% neutral buffered formalin. Tissues were embedded in paraffin for tissue section (2 μm thickness) preparation and Hematoxylin and eosin (H&E) staining at the University of Vermont Microscopy Imaging Center and University of Vermont Medical Center.

The Omicron BA.2 virus challenge study was carried out at Bioqual Inc. (Rockville, MD) in compliance with local, state, and federal regulations with approval by Bioqual and Institutional Animal Care and Use Committees (IACUC). In brief, 6–8 weeks old male K18-hACE2 transgenic mice (Jackson Laboratory) were randomly assigned to different challenge groups, anesthetized with a cocktail of Ketamine/Xylazine/antisedan (80mpk/5mpk/1mpk) and then inoculated intranasally with 7Χ10^3^ TCID_50_ of Omicron BA.2 virus preparation (NR-56522, lot#70051594) 6 hour before or after intranasal instillation with purified mACE2-IgM ranging from 0.625 to 5mg/kg or the control (a nonrelated IgM or phosphate-buffered saline, PBS). Morbidity was monitored by body weight loss after infection and clinical signs of infection including hunching, ruffling of fur, lethargy, respiratory distress, and death daily for a total of 14 days. The percentage body weight was determined relative to the starting weight before viral challenge. When losing greater than 20% body weight, mice were humanely euthanized and considered dead. On day 4 post-challenge, mice were humanely euthanized from each group and lung samples removed and quickly frozen in liquid nitrogen for viral load titration.

### Lung viral titration

For WT WA1 viral titration, weighted lung tissue was thawed and then bead beat in 750 μL of Dulbecco’s minimum essential medium (DMEM, Gibco, 11965-092) with 1% cocktail of penicillin and streptomycin (Gibco, 15140-122). Ten-fold dilutions of lung homogenate were assayed in duplicate (100 μL of diluted sample per well) and allowed to adsorb for 1 hour on Vero E6 cells. The inoculums were then washed off and overlaid with approximately 500 μL of a 1:1 mix of 2X DMEM and 1.4% agarose. After 3 days, plates were fixed with the addition of 25% formaldehyde and monolayers stained with 1% crystal violet. Counted plaques were normalized to total tissue mass for calculations of plaque forming units per gram of lung tissue. For Omicron BA.2 viral titration, lungs were weighed, homogenized in cell culture medium, centrifuged and the resulting supernatants were used to determine viral titer by inoculating serial dilutions of samples on Vero-TMPRSS2 cells. Infected plates were incubated at 37°C with 5% CO2 for four days before the wells were visually inspected for cytopathic lesions. TCID_50_ of all samples were calculated using the Read-Muench formula and reported as TCID_50_ per gram of lung.

### Aerosolization

Aerosolization of mACE2-IgM was performed using a Teleflex® MAD300 atomizer or an Aerogen® Solo vibrating mesh nebulizer. Evaluation of atomization based aerosolization was conducted by drawing 0.3 mL of 1 mg/mL drug product into a sterile disposable silicone oil-free syringe, attaching the atomizer, and hand actuating solution into a sterile collection tube aseptically within a class II biosafety cabinet. Similarly, evaluation of nebulization based aerosolization was conducted by directly filling 0.5 mL of approximately 1 mg/mL drug product into the nebulizer within a class II biosafety cabinet. Drug product was nebulized at a fixed rate into a sealed sterile collection tube within a class II biosafety cabinet. Visual observation and dose duration were recorded during collection to evaluate device performance. The aerosolized mACE2-IgM was stored at 37°C, 25°C, 4°C and −70°C for stability monitoring via SEC analysis and ELISA assay.

### Statistical analysis

All statistical analyses were performed using GraphPad Prism 9 and the statistic tests are described in the indicated figure legends. Nonlinear regression curve fitting was used for calculating the EC_50_ and IC_50_ values. P value equal or less than 0.05 was considered statistically significant.

## Supplementary Material

**Fig. S1.**
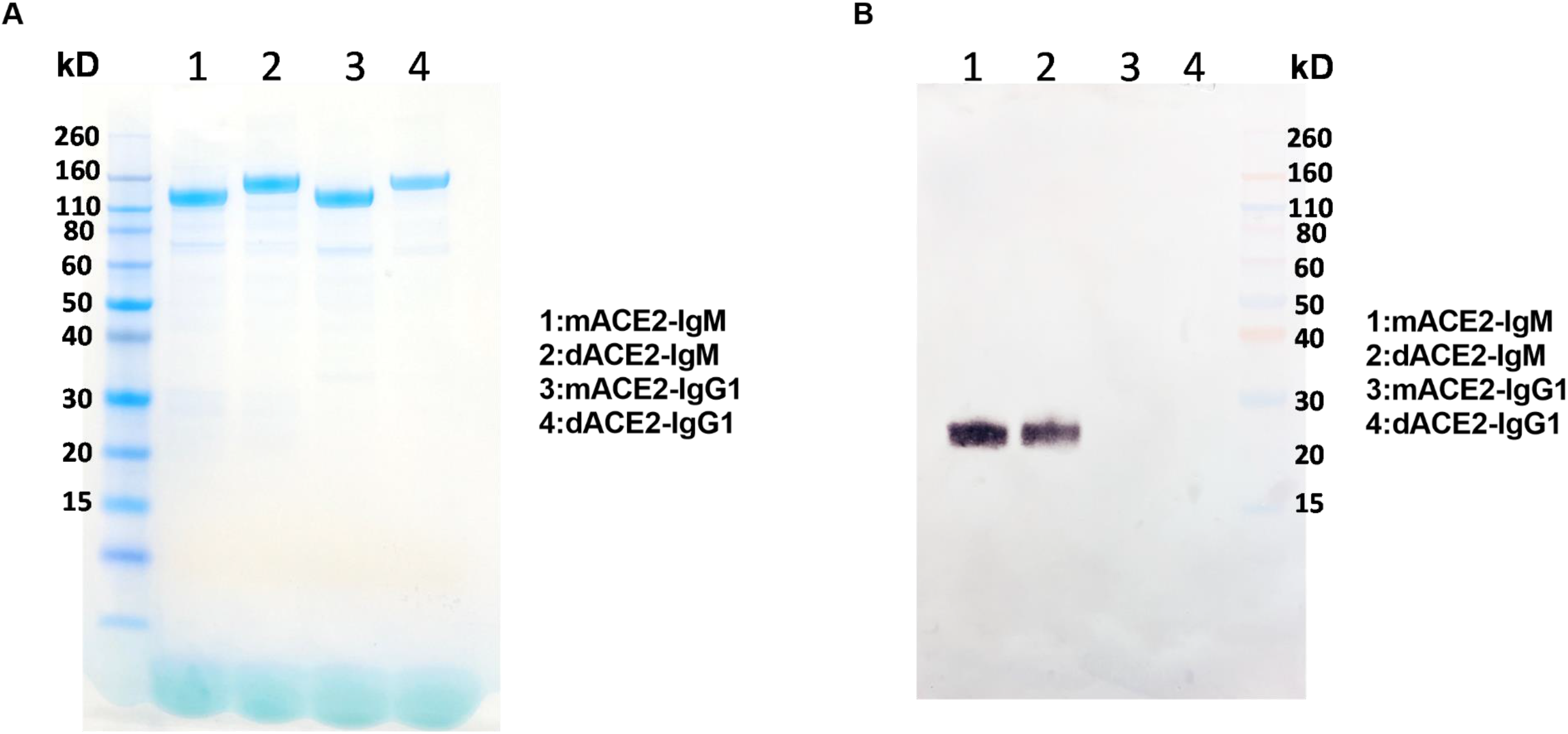
Biochemical analysis of ACE2 constructs. (**A**) Reducing SDS-PAGE gel of mACE2-IgM, dACE2-IgM, mACE2-IgG1, and dACE2-IgG1 with Coomassie blue staining. (**B**) Western Blot of mACE2-IgM, dACE2-IgM, mACE2-IgG1, and dACE2-IgG1 using anti-J chain antibody.

**Fig. S2.**
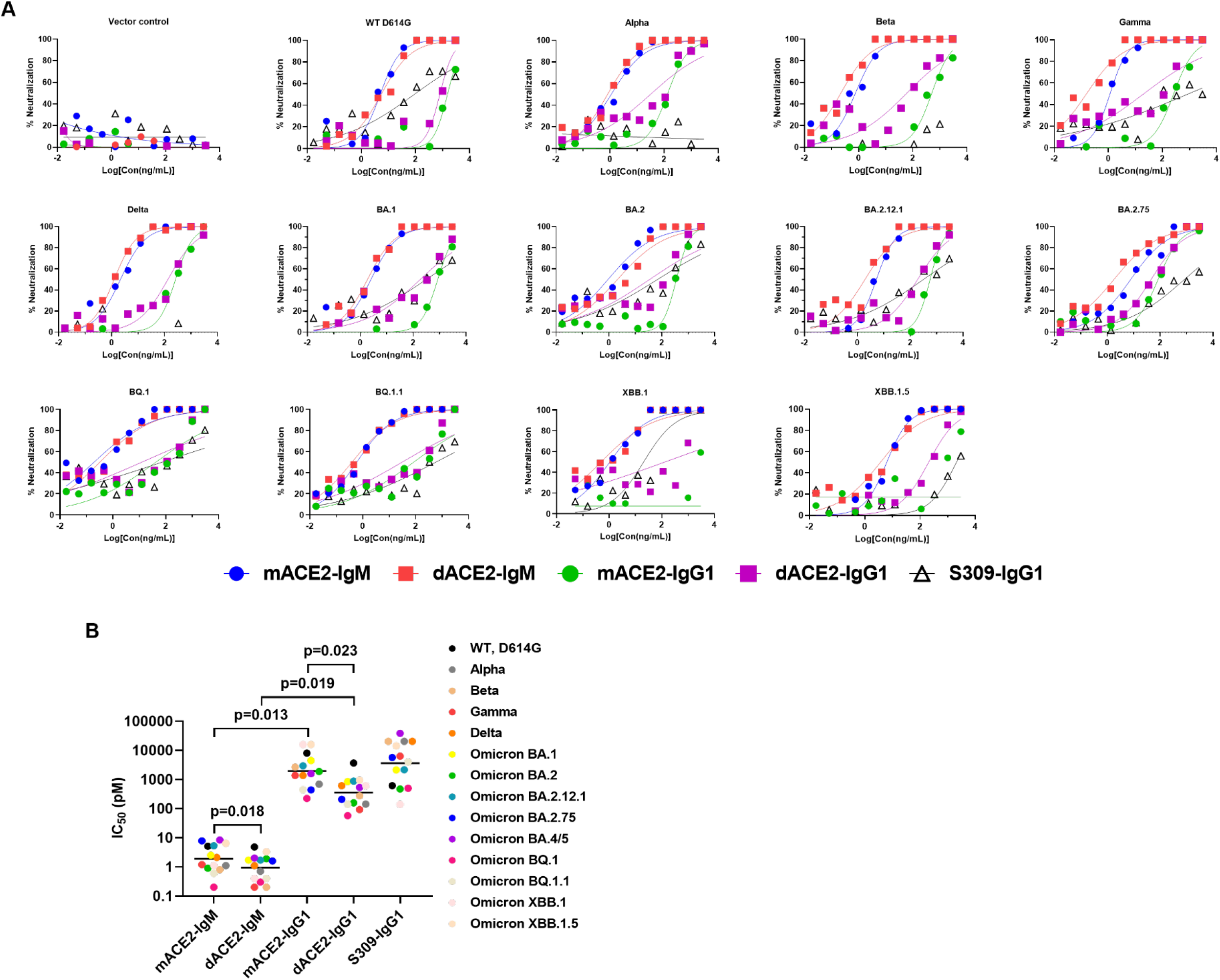
Virus neutralization. (**A**) A panel of pseudoviruses containing spike proteins of different SARS-CoV-2 variants and SARS-CoV-1 virus were tested against mACE2-IgM, dACE2-IgM, mACE2-IgG1, dACE2-IgG1 and control S309 IgG1. IC_50_ values were calculated based on four parameter curve fitting. (**B**) IC_50_ (pM) of mACE2 and dACE2 decamer against a panel of pseudotyped SARS-CoV-2 variants and SARS-CoV-1 virus. The panel includes major variants of Alpha, Beta, Gamma, Delta, and Omicron BA.1, BA.2, BA.4/5, BQ.1, BQ.1.1, XBB.1, and XBB.1.5, and each symbol represents one individual variant. The horizontal line indicates the geometric mean for the cluster. The IC_50_ (pM) values are listed in table S2. Statistical differences between indicated groups were analyzed by two-tailed paired t-test.

**Fig. S3.**
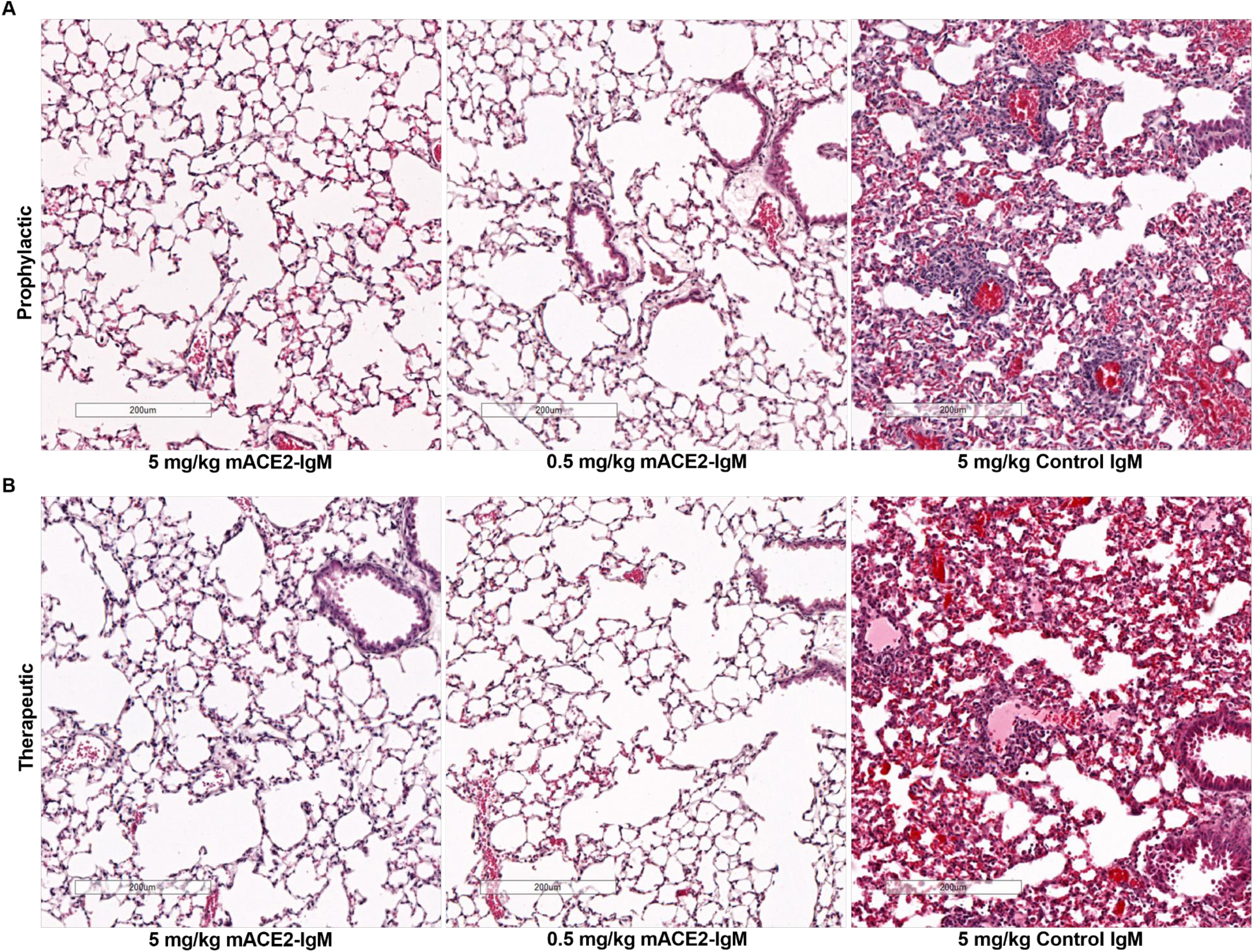
Histopathological analysis of SARS-CoV-2 infection. K18 hACE2 transgenic mice were treated intranasally (12 per group) with mACE2-IgM or control IgM 6h before or after intranasal challenge with 1000 PFU dose of WT WA1 virus. Hematoxylin and eosin (H&E) staining was performed on the lung tissues collected at day 3 after viral infection in mice pre-treated (A), or post-treated (B) with 5 mg/kg, 0.5 mg/kg of ACE2-IgM or 5 mg/kg control IgM. Scale bars, 200 μm. Images are representative of n=4 per group.

**Fig. S4.**
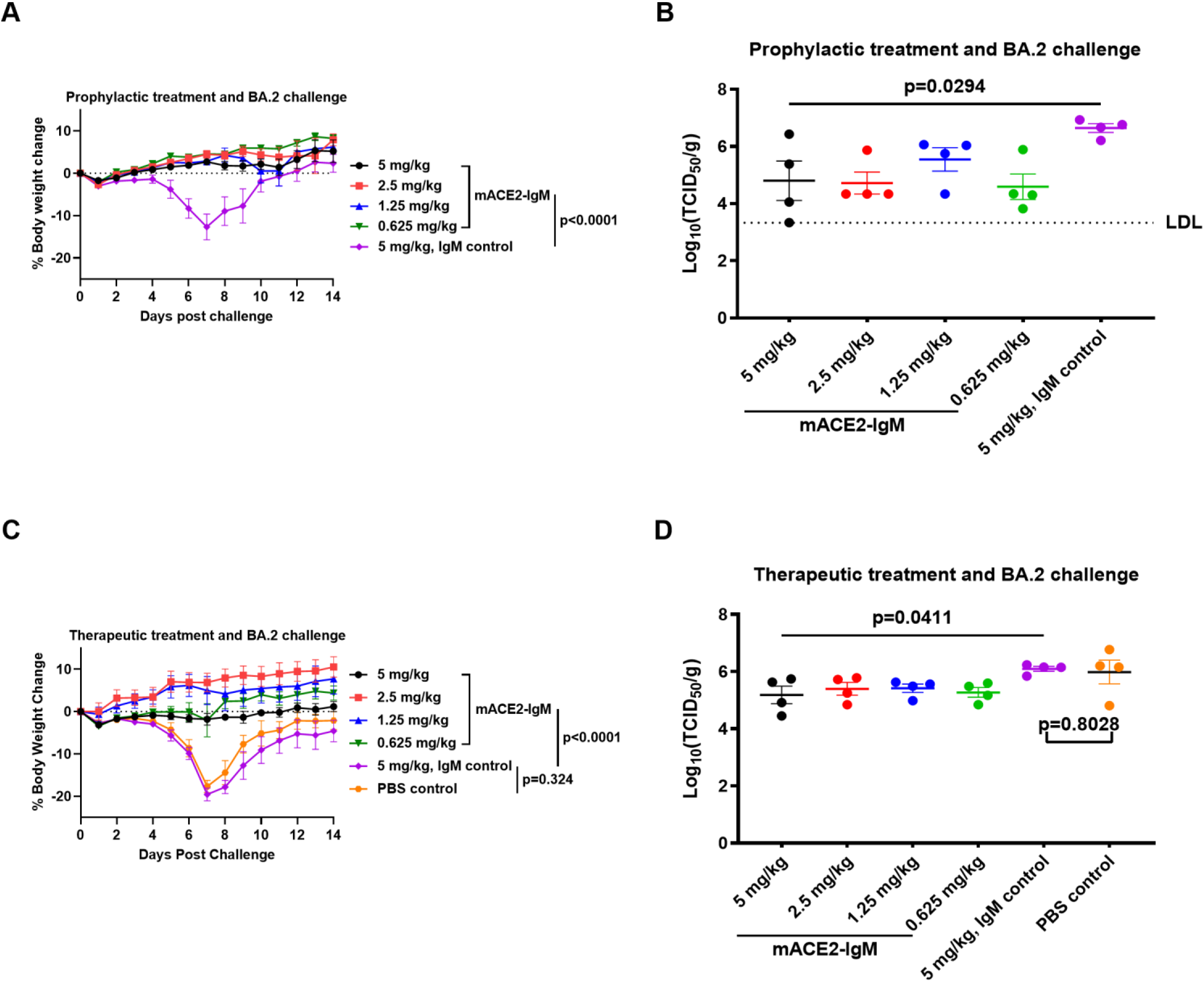
In vivo protection of ACE2 decamer. K18 hACE2 transgenic mice were treated intranasally (10-12 per group) 6h before or after intranasal challenge with 7.0X10^3^ TCID_50_ of Omicron BA.2 virus. (**B**) Weight change and (**C**) viral titer of lungs of animals treated intranasally with mACE2-IgM or control 6h before intranasal viral challenge. (**D**) Weight change and (**E**) viral titer of lungs of animals treated with mACE2-IgM or control 6h after intranasal viral challenge. The data are means ± SEM for weight change and viral load in each group. Two-way ANOVA (mixed-effects model) was used for comparisons of weight change. Ordinary one-way ANOVA was used for comparison of lung viral titers of multiple groups and two-tailed unpaired t-test was used for comparison of IgM and PBS control group. LDL: lower detection limit.

**Fig. S5.**
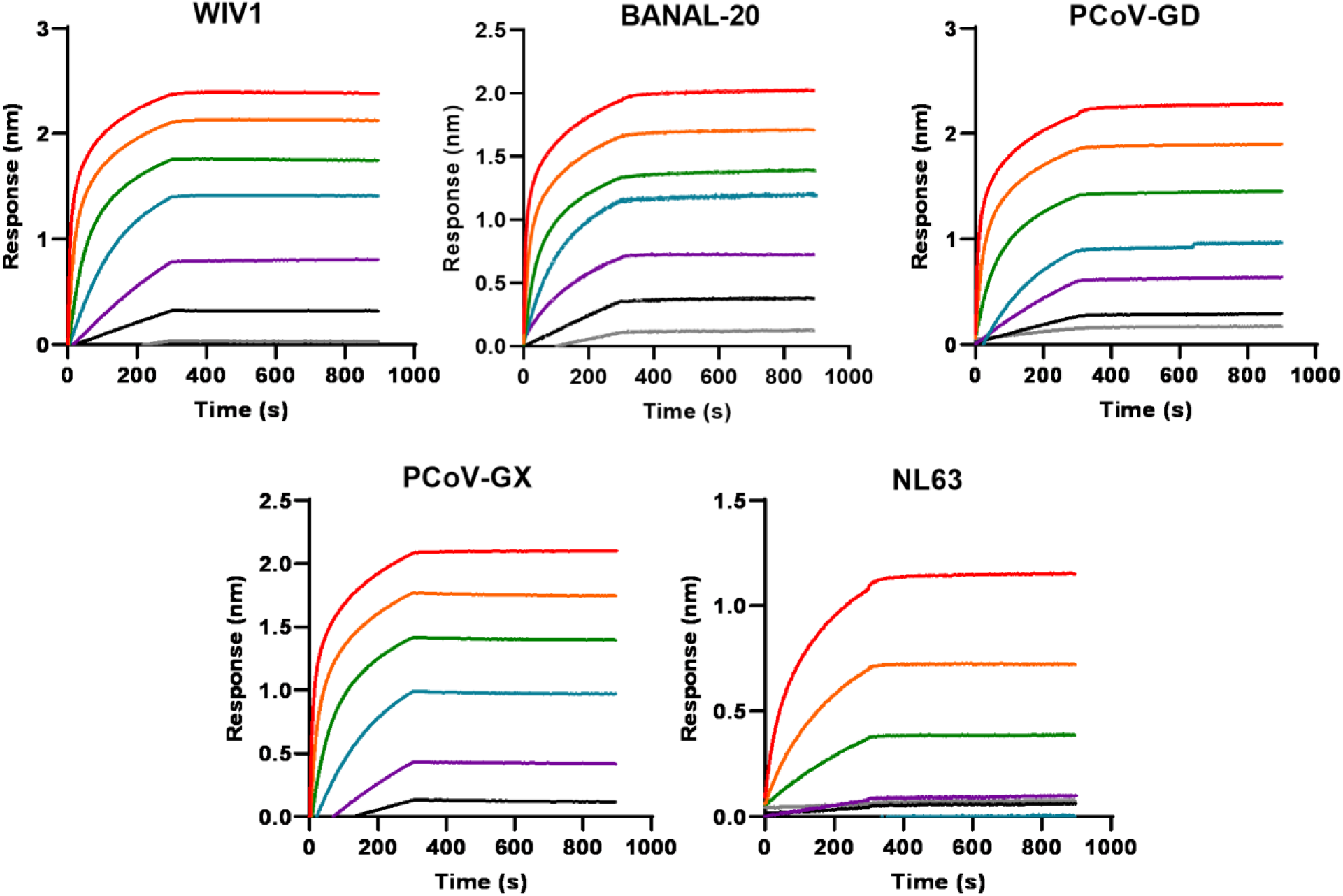
Binding of RBDs of animal and NL63 CoV. Biolayer interferometry analysis of the binding with mACE2-IgM decamer by RBDs of WIV-1, BANAL-20, PCoV-GD, PCoV-GX, and NL63 viruses. ForteBio Octet data analysis software was used to calculate the *K*_D_ data with the global fitting method.

**Table S1:** Biolayer interferometry RBD binding by ACE2-IgM.

| Molecule | $K_a(1/Ms)$ | $K_{dis}(1/s)$ | $K_D(M)$ |
| --- | --- | --- | --- |
| **mACE2-IgM** | 9.5E+04 | <1.0E-07 | <1.0E-12 |
| **dACE2-IgM** | 1.1E+05 | <1.0E-07 | <1.0E-12 |
| **mACE2-IgG1** | 1.6E+04 | 3.5E-04 | 2.2E-08 |
| **dACE2-IgG1** | 1.8E+04 | 2.7E-04 | 1.5E-08 |

**Table S2:** IC_50_ summary of pseudovirus neutralization assay for SARS-CoV-2 WT and variants.

|  | IC<sub>50</sub>(ng/mL) |  |  |  |  |
| --- | --- | --- | --- | --- | --- |
|  | mACE2-IgM | dACE2-IgM | mACE2-IgG1 | dACE2-IgG1 | S309-IgG1 |
| WT | 5.0 | 5.4 | 1526.0 | 795.8 | 89.9 |
| Alpha | 1.1 | 0.8 | 132.0 | 31.2 | >3000 |
| Beta | 0.8 | 0.2 | 506.4 | 60.0 | >3000 |
| Gamma | 1.2 | 0.2 | 260.8 | 20.3 | 922.8 |
| Delta | 2.1 | 1.2 | 262.4 | 134.0 | >3000 |
| BA.1 | 2.5 | 1.9 | 852.3 | 185.5 | 312.7 |
| BA.2 | 0.9 | 2.1 | 356.1 | 34.7 | 69.1 |
| BA.2.12.1 | 5.2 | 1.9 | 563.6 | 191.0 | 316.7 |
| BA.2.75 | 7.7 | 1.8 | 84.1 | 45.7 | 832.3 |
| BA.4/5 | 8.3 | 2.3 | 299.3 | 116.8 | 5648.0 |
| BQ.1 | 0.2 | 0.3 | 42.4 | 12.5 | 73.4 |
| BQ.1.1 | 0.6 | 0.4 | 84.8 | 30.9 | 581.6 |
| XBB.1 | 1.1 | 0.5 | >3000 | 133.5 | 20.4 |
| XBB.1.5 | 6.2 | 3.7 | >3000 | 205.1 | 2087 |

|  | IC<sub>50</sub>(pM) |  |  |  |  |
| --- | --- | --- | --- | --- | --- |
|  | mACE2-IgM | dACE2-IgM | mACE2-IgG1 | dACE2-IgG1 | S309-IgG1 |
| WT | 5.1 | 4.8 | 7989.5 | 3633.8 | 611.6 |
| Alpha | 1.1 | 0.7 | 691.0 | 142.5 | >20408 |
| Beta | 0.8 | 0.2 | 2651.3 | 274.0 | >20408 |
| Gamma | 1.2 | 0.2 | 1365.4 | 92.7 | 6277.6 |
| Delta | 2.1 | 1.1 | 1373.8 | 611.9 | >20408 |
| BA.1 | 2.5 | 1.7 | 4462.3 | 847.0 | 2127.2 |
| BA.2 | 0.9 | 1.9 | 1864.4 | 158.4 | 470.1 |
| BA.2.12.1 | 5.3 | 1.7 | 2950.8 | 872.1 | 2154.4 |
| BA.2.75 | 7.8 | 1.6 | 440.3 | 208.7 | 5661.9 |
| BA.4/5 | 8.4 | 2.0 | 1567.0 | 533.3 | 38421.8 |
| BQ.1 | 0.2 | 0.3 | 222.0 | 57.1 | 499.3 |
| **BQ.1.1** | 0.6 | 0.4 | 444.0 | 141.1 | 3956.5 |
| **XBB.1** | 1.1 | 0.4 | >15706.8 | 609.6 | 138.8 |
| **XBB.1.5** | 6.4 | 3.3 | >15706.8 | 936.5 | 14197.3 |

**Table S3:** Mass recovery, structural integrity, and functional analysis of mACE2-IgM upon atomization and nebulization.

|  | %Mass Recovery<sup>†</sup> | %HMW<sup>‡</sup> | %Main<sup>§</sup> | %LMW<sup>§</sup> | %Binding<sup>*</sup> |
| --- | --- | --- | --- | --- | --- |
| **Pre-aerosolization** | 100 | 0.5 | 93.1 | 6.4 | 100 |
| **Post atomization** | 100 | 0.7 | 93.6 | 5.7 | 112 |
| **Post nebulization** | 100 | 0.5 | 99.1 | 0.4 | 101 |

<sup>†</sup>Mass recovery was determined by SEC-MALS analysis.
<sup>‡</sup>HMW: high molecular weight.
<sup>§</sup>Main is for the major peak in the SEC chromatogram.
<sup>§</sup>LMW: low molecular weight.
<sup>\*</sup>Binding was determined by ELISA.

## Acknowledgement

We appreciate the scientists and animal care takers at Bioqual, Inc for coordinating and conducting the Omicron BA.2 challenge study at its BSL3 animal facility and we also would like to thank the scientists of NanoImaging Services for performing the TEM study. This work was funded by IGM Biosciences, Inc. Work in Dr. An’s laboratory was funded in part by the Welch Foundation grant no. AU-0042-20030616.

## Author contributions

TF, PRH and HG conceived the concept and designed constructs. KX contributed to the initial structural analysis. BC, YY, SJH, SH, MD, KC, SH, SS, CM, AF, RH, AW, PE, AF, RH, AKR and JPS conducted experiments and generated critical data. NZ, ZA, JWB, SFC, BAK, AWW, WRS, and JWS contributed to discussion and program strategies. HG coordinated and managed the projects. HG and TF wrote the manuscript. All authors read and approved the final manuscript.

## Data and materials availability

All source data associated with this study present in the paper, or the Supplementary Materials are available upon request under confidentiality agreement. Further data can be obtained by contacting the corresponding authors.

